# LSD-pipeline: Causal Inference of miRNA Network Effects in Alzheimer’s Disease

**DOI:** 10.64898/2026.09.17.747423

**Authors:** Jian Shi

## Abstract

MicroRNAs (miRNAs) are implicated in Alzheimer’s disease (AD), but research has focused on individual miRNAs and direct targets. Existing approaches to miRNA regulation in AD identify associations rather than causal effects, and few methods estimate multi-stage chains from miRNAs through target genes to target transcription factor (TF) cascades. We developed the LSD pipeline (LASSO-SEM-DoWhy), integrating LASSO feature selection, multi-stage structural equation modeling, and DoWhy causal inference to identify and validate miRNA causal pathways in AD. Applying LSD to six blood miRNA and brain mRNA datasets, we identified four LSD-validated miRNAs (miR-30d-5p, miR-92a-3p, miR-296-5p, miR-193a-5p) as AD biomarkers, achieving >86% ROC accuracy in an independent validation cohort. Several miRNAs with no significant direct association with AD showed significant effects when estimated through their target networks, while others significant in direct analysis were not supported at the network level, underscoring the value of network-level analysis. Extending to the TF layer revealed complete miRNA → targets → TF cascades → AD causal chains, with HMGA1, NKX2-3, and PRRX2 as key intermediaries. Confirmed classic pathways converge primarily on tau pathology and synaptic dysfunction. miRNA effects were largely age-independent, suggesting miRNAs act as early initiators of AD pathogenesis. Beyond AD, the LSD pipeline provides a generalizable framework for uncovering causal regulatory mechanisms in other diseases.

## Introduction

Alzheimer’s disease (AD) is an incurable condition that affects approximately 50 million people worldwide, with the incidence rate continuing to rise each year ^[1]^. The classic pathological mechanisms of AD include β-amyloid (Aβ) plaques, neurofibrillary tangles composed of hyperphosphorylated tau proteins, and progressive neurodegeneration. As AD progresses, it becomes increasingly burdensome for patients, their families, and society as a whole. Early detection can help slow or delay the disease’s progression; however, the initial mechanisms remain unclear. Recently, causal inference and multi-omics approaches, including those applied to miRNAs, have begun to reveal the initial and causal mechanisms of AD.

Although miRNAs regulate gene expression through extensive target networks, most studies focus on single miRNA-target interactions, which cannot capture effects that emerge only at the network level. This narrow approach is inadequate because a single miRNA can target tens to hundreds of genes, while individual genes are typically regulated by multiple miRNAs [2]. However, the regulatory mechanisms of these networks are not yet clear and remain largely unexplored. MiRNAs, which are 20-22 nt non-coding RNAs, contribute to the development, function, and progression of neurodegenerative diseases, including AD by negatively regulating mRNA expression by binding to the non-coding 3’ UTRs of targeted mRNAs [3–7]. To capture these network-level effects on AD, we move beyond descriptive associations and adopt counterfactual reasoning to investigate how individual miRNAs regulate their targets within the broader network. The potential-outcomes framework provides a conceptual basis for defining the causal effects of multiple miRNAs and their target genes.

Currently, miRNAs are widely used as biomarker candidates in clinical research, measured in blood (circulating miRNAs) and other body fluids [8–10]. Evidence is accumulating that miRNA levels in human body fluids (which may originate from the brain or other organs) are associated and reflected with the development and progression of AD, as well as other diseases [7, 10–13], as illustrated in Fig 1A. This suggests that circulating miRNAs can serve as both potential biomarkers for AD diagnosis and sources of insight into AD pathogenesis. However, current approaches for selecting miRNA biomarkers predominantly rely on statistical methods, without incorporating biological or causal functions related to their origination. In this study, we identify miRNA biomarkers through causal analysis: estimating the effect of an exposure (miRNA expression) on the outcome (AD status) under an assumed causal structure, with explicit assumptions and refutation testing, not measuring statistical association. This yields candidates that demonstrate both statistical robustness and mechanistic relevance to AD pathogenesis.

**Figure 1.**
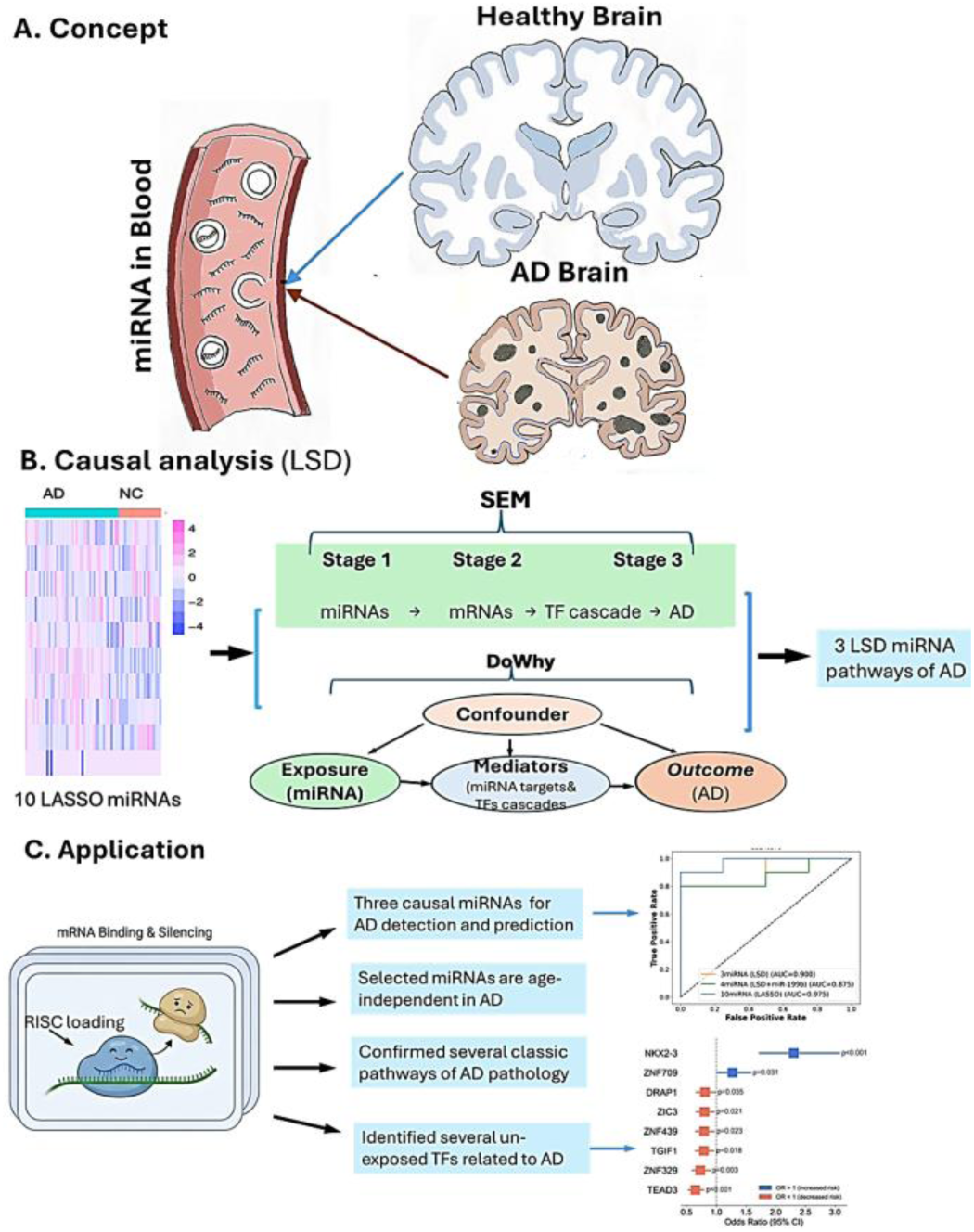
The LSD Framework of Causal Analysis for AD. A) Concept: Serum miRNAs may originate in the brain and exert causal effects on target genes within the brain, including TFs and their downstream cascades, thereby reflecting the onset and progression of AD. B) Causal analysis: In the LSD pipeline we developed, LASSO is used for feature preselection; SEM is used for multi-stage causal exploration (miRNAs → AD, mRNAs → AD, TF targets → AD); and DoWhy validates the causal effects identified by SEM through DAG-based causal inference, explicitly adjusting for confounding factors, including age. C) Application: LSD-identified causal miRNAs that are independent of age were used to detect AD using ML/DL models. Functional analysis of significant miRNA pathways revealed several TFs and their downstream cascades as potential drivers of AD, as well as classic AD pathology.

Causal analysis is a powerful method for investigating disease mechanisms, but its application to miRNA studies in AD has been limited by stringent data requirements. These requirements include either variables from a single source [14] or genomic variation data such as SNPs and their related factors [15, 16]. We propose to overcome these limitations by leveraging a unique property of miRNAs: they regulate their target genes through highly conserved binding sites and seed sequences, which are shared even across species [17]. This biological stability and target-binding specificity facilitate robust causal inference from multi-source data.

Building on this insight, we developed the LSD pipeline (LASSO–SEM–DoWhy) [18] to systematically identify and validate miRNA causal pathways in AD, as shown in the middle panel of Fig. 1B. The pipeline comprises three sequential components: LASSO feature selection, multi-stage structural equation modeling (SEM) [19], and DoWhy validation. The SEM component, implemented in lavaan, is organized into three nested stages: stage 1 estimates miRNA effects on AD; stage 2 estimates the effects of mRNA targets; and stage 3 estimates the effects of TF target genes. Subsequently, at the pipeline level, DoWhy [20] normalizes the assumed structure as a directed acyclic graph (DAG) and applies refutation tests, which include placebo treatments, random common causes, and subset exclusion, to assess the robustness of the estimated effect against violations of identification assumptions. This involves specifying DAGs with explicit exposure (miRNA), mediators (miRNA targets and transcription factor (TF) cascades), outcome (AD), and confounder (age), as shown in the lower section of Fig. 1B).

Our primary methodological contribution is the LSD pipeline [18], designed for studying the causal effects of miRNAs in identifying LSD miRNAs as biologically significant biomarker candidates and assessing their predictive performance using various machine learning (ML) and deep learning (DL) models (Fig. 1C). Notably, these LSD miRNAs are largely independent of age, making them promising candidates for the early prediction and detection of AD. Functional analyses (including Gene Ontology (GO), Kyoto Encyclopedia of Genes and Genomes (KEGG), WikiPathway (WP), and Reactome (REAC)) of the significant miRNA pathways confirm several established AD pathological pathways. Additionally, they suggest that the dysregulation of TF cascades driven by miRNAs may act as potential contributors to AD pathogenesis (Fig. 1C). Overall, these findings underscore the effectiveness of the LSD pipeline in revealing miRNA-specific causal mechanisms of AD, which are challenging to capture with traditional methodologies, while also highlighting its potential applicability to other diseases.

## Methods

### Data collection and web databases and tools

This study included datasets from miRNA and mRNA studies. The study was divided into two phases: the discovery phase and the validation phase. During the discovery phase, following data cleaning, this study utilized miRNA expression data derived from the GSE120584 [21] blood sample dataset, which comprised 980 AD, 32 mild cognitive impairment (MCI), and 279 normal control (NC) samples. In a mRNA dataset for discovery, GSE33000 included 310 AD and 157 non-dementia (ND) brain (prefrontal cortex) tissue samples [22]. In total, 1,290 AD, 32 MCI, 279 NC, and 157 ND were included in this phase. For the validation phase, two miRNA sequence datasets of GSE215789 [23] and GSE46579 [24] from blood were combined with 76 AD, 62 MCI and 43 control samples. For mRNA datasets for validation, GSE36980 [25] dataset with 33 AD and 47 control samples from front cortex (prefrontal cortex) tissues were combined with GSE122063 [26] dataset with 12 AD and 11 control samples also from FC tissues.

The web data bases for miRNA target gene and TF analyses included TargetScan 8.0 (https://www.targetscan.org/vert_80/), Transcriptional Regulatory Relationships Unraveled by Sentence-based Text mining (TRRUST) human dataset (https://www.grnpedia.org/trrust/), CollecTRI (https://github.com/saezlab/CollecTRI) [27], and DoRothEA (v3.22) (https://dorothea.opentargets.io/). For functional analyses, the databases included Gene Ontology (GO) (http://geneontology.org/) (released 8-5-2026), Kyoto Encyclopedia of Genes and Genomes (KEGG) (https://www.genome.jp/kegg/pathway.html, Release 119.0), Reactome (https://reactome.org/, Release 1.6.8), WikiPathways (https://www.wikipathways.org/) (PMID: 37941138) and g:Profiler (version e113_eg59_p19).

All datasets were obtained from the Gene Expression Omnibus (GEO) and consist of publicly available, de-identified samples; the original data collection was approved by the contributing institutions’ IRBs, and secondary analysis of these data was exempt from additional IRB review.

All gene and transcription factor symbols follow HGNC nomenclature; official gene names can be retrieved from the HGNC database (https://www.genenames.org).

### Differential Expression Analysis

Jonckheere-Terpstra (JT) trend (clinfun (1.1.5) in R) analysis is a non-parametric, rank-based statistical method used to detect ordered trends (increasing or decreasing) across three or more independent, ordered groups. The miRNA dataset includes NC, MCI, and AD samples, and JT trend analysis was applied to determine whether miRNA expression levels increase or decrease monotonically across the ordered progression of AD stages. For mRNA data, differential expressions between AD and ND brain tissue samples were assessed using an independent samples Student’s t-test, as only two groups were compared. Given the exploratory nature of the discovery stage, significance thresholds were set at p<0.05 for both analyses. FDR-adjusted q-values (Benjamini-Hochberg procedure) and p-values are reported for both the Jonckheere-Terpstra trend analysis (miRNA) and Student’s t-test. Statistical details will be provided on each table, figure, and supplement. Specifically, at stages 2–3, the analysis is restricted to predicted targets of the stage 1 miRNAs rather than a genome-wide scan. The priority at these stages is sensitivity: excluding true targets truncates the causal chain and biases downstream estimates toward the null. We therefore maintained p < 0.05 at these stages, accepting a higher false-positive rate in exchange for broader coverage.

### Target Gene Selection

The miRNA candidates and their target gene candidates were selected and identified using TargetScan (v8.0) and miRDB v6.0 (https://mirdb.org/) [28]. TS conserved-site prediction file contained 313 unique human miRNAs with evolutionarily conserved predicted binding sites, while miRDB identified over 2,600 known human miRNAs in total. There are 271 miRNAs overlapping between these databases, and this study focused on these miRNAs. To retain only high-conservative miRNAs, we applied a context++ score threshold of ≤ −0.50 in TS and a score ≥ 80 in miRDB. The miRNA target gene candidates were selected from TS with the same context++score. Some unique genes with multiple test probes were averaged to maintain a singular position in the target list. This composite metric integrates site type, 3’ compensatory pairing, local AU content, target site accessibility, and evolutionary conservation [29]. This threshold was chosen a priori based on empirical evidence indicating that more negative context++ scores correspond to greater target repression efficacy. Among the filtered target genes, transcription factors (TFs) were identified using TRRUST [30] (https://www.grnpedia.org/trrust/), and their downstream targets were obtained from the CollecTRI [27] and DoRothEA [31] databases.

### LASSO-based miRNA feature selection

Based on significantly miRNA expressed data from JT analysis, these miRNAs were mapped from TargetScan, and then, the TS selected targeted genes were mapped with significant mRNA data. Subsequent LASSO feature selection provided an additional layer of logistic regression, refining the candidate set to a small subset (Fig. 1B), thereby mitigating concerns about multiple testing. We compared LASSO logistic regression and LASSO regularization regression. Considering the number of target genes per miRNA and the requirements of downstream SEM analysis, LASSO logistic regression was selected for this study. The R package glmnet (v 5.0) was used to implement LASSO logistic regression with L1 penalty, optimized via 5-fold cross-validation using the saga solver. LASSO feature selection, SEM causal analysis, and DoWhy validation have been integrated into the LSD package [18] (https://doi.org/10.5281/zenodo.20369677)).

### Causal Pathway Analysis Using Structural Equation Modeling (SEM)

#### Structural Equation Modeling Framework

SEM analysis was conducted using the lavaan package (v0.6-16) in R 4.2.2 [32]. Based on the regulatory hierarchy among miRNA targets, we employed either a two-stage model for non-transcription factor targets or a three-stage model when transcription factor intermediates were identified, as shown in the middle panel of Fig. 1B. The two-stage model characterized the pathway as miRNA→target gene→AD, while the three-stage model extended this to miRNA→TF→TF target gene→AD. Since miRNA and mRNA expression data were derived from separate patient cohorts, each stage was estimated independently, and the results were integrated to reconstruct complete causal pathways.

#### Stage 1: miRNA → AD Status

Twenty miRNAs selected via LASSO were modeled as predictors of AD diagnostic status. Given the binary nature of the outcome, a probit model was specified: Y = βᵢXᵢ + ε, i = 1, 2, …, 20, where Y is the observed binary AD status (0 = NC, 1 = AD), Xᵢ represents the expression level of the i-th miRNA (i = 1, …, 20), βᵢ is the corresponding path coefficient, and ε is the error term. The model was estimated using the weighted least squares mean and variance adjusted (WLSMV) estimator, which is appropriate for categorical endogenous variables [33]. miRNAs with significant path coefficients (p < 0.05) were carried forward to integration. The impact of these miRNAs on AD risk is dependent on β values: if β > 0, the miRNA increases AD risk; if β < 0, it decreases AD risk.

#### Stage 2: Target Genes → AD Status

All filtered target genes of LASSO-selected miRNAs were modeled as predictors of AD status in Stage 2. Stage 2 focuses on miRNA-level causal effects mediated through downstream target networks rather than individual gene-level effects. All Stage 2 models were estimated by using maximum likelihood robust (MLR). The modeling approach depended on the number of available target genes and non-TF targets or TF targets for each miRNA:

a) Non-TF target genes: When fewer than three target genes were available, stage 2 analysis was excluded because of insufficient indicators for latent factor identification. When the three to five target genes were available, a latent factor model was specified:

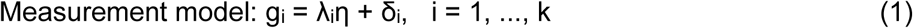

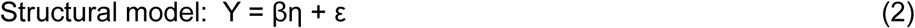

where g_i_ are observed target gene expression levels, λ_ᵢ_ are factor loading (gene’s contribution to latent construct), k is the number of available target genes (3≤ k <6), β are path coefficients, and ε is the error term.

For miRNAs with ≥ 6 non-TF target genes, two complementary latent factor modeling approaches were estimated for each miRNA:

Latent model was same as above using formula (1) and (2) to represent the miRNA’s downstream regulatory effect, in which k is the number of target genes. After that, the latent factor model was fitted; if model fit was poor (CFI < 0.85, confirmed by RMSEA > 0.10 and SRMR > 0.10), item-to-construct balance parcel model was applied as below.

Parceled model was reserved for k ≥ 6 because with fewer indicators, parcels would contain too few genes to meaningfully reduce measurement error. In this model, target genes were combined into several parcels using item-to-construct balance parceling (see Item-to-Construct Balance Parceling below).

b) TF target genes: For miRNAs with fewer than three TF targets, a direct observed variable regression was specified:

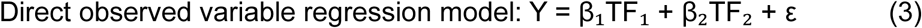

where TF_₁_ and TF_₂_ are observed TF expression levels, β_₁_ and β_₂_ are individual path coefficients, and ε is the error term.

For TF targets ≥ 3, the same latent model of formula (1) and (2) was used. No parceling applied in Stage 2 for TFs as miRNA targets.

### Stage 3: TF Target Genes → AD Status (3-Stage Pathway Only)

For miRNA target genes identified as TFs with known downstream targets in CollecTRI or DoRothEA, an additional modeling stage (Stage 3) was conducted to capture individual TF-level effects within miRNA–TF cascades. The modeling approach depended on the number of available downstream target genes per TF: when fewer than 3 downstream target genes were available, a direct observed variable regression model was specified using formula (3); when 3–5 downstream target genes were available, a latent factor model was fitted using formulas (1) and (2) without parceling; when ≥ 6 downstream target genes were available, a latent factor model was fitted first using formulas (1) and (2), and model fit was evaluated using the same criteria as above (CFI, RMSEA, SRMR). If model fit was poor, item-to-construct balance parceling was applied as described below.

#### Item-to-Construct Balance Parceling

For latent factor models with poor fit (CFI < 0.85, confirmed by RMSEA > 0.10 and SRMR > 0.10), item-to-construct balance parceling was implemented to reduce measurement error and improve indicator reliability [32, 34]. Models with fewer than 6 target genes were ineligible for parceling. The number of parcels was determined dynamically as:

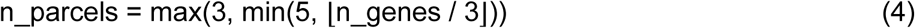

Target genes were rank-ordered by the magnitude of their standardized factor loadings and distributed into parcels using a snake-order assignment to ensure each parcel contained a balanced mix of genes with strong and weak loadings. For example, with 3 parcels: Parcel 1 received genes ranked 1st, 6th, 7th, …; Parcel 2 received genes ranked 2nd, 5th, 8th, …; and Parcel 3 received genes ranked 3rd, 4th, 9th, …. Parcel scores were calculated as the means of standardized expression values (z-scores) of the constituent genes. The latent factor model was then re-estimated using m parcels as indicators:

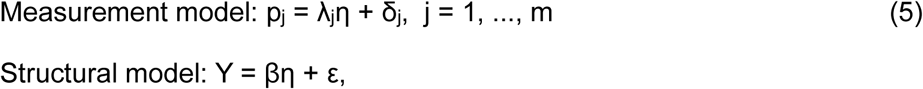

where p_₁_, …, p_ₘ_ are parcel scores, m is the number of parcels (3 ≤ m ≤ 5), η is the latent regulatory factor representing the miRNA’s downstream effect, and all other terms are as defined above.

#### Model Fit Assessment

Model fit was evaluated using three complementary indices: the Comparative Fit Index (CFI; ≥0.95 indicates good fit, ≥0.90 indicates acceptable fit) [35], the Root Mean Square Error of Approximation (RMSEA; <0.06 indicates good fit, <0.08 indicates acceptable fit), and the Standardized Root Mean Square Residual (SRMR; <0.08 indicates acceptable fit) [35]. Models that did not achieve acceptable fit after parceling were reported as negative findings.

#### Parameter Estimation and Integration

All path coefficients are reported as standardized values (β). Stage 1 models were estimated by the WLSMV estimator to accommodate the binary AD outcome. Stage 2 and Stage 3 models, however, were estimated with MLR [36] in lavaan. Standardized path coefficients (β), their standard errors, and p-values were extracted from the lavaan output, with paths having p < 0.05 considered statistically significant. For latent-factor models, standardized factor loadings (λ) were examined to assess the contribution of individual genes to the latent construct. Significant paths were then chained across stages to reconstruct complete causal pathways: miRNA → target gene → AD (two-stage model) or miRNA → TF → TF-target gene → AD (three-stage model). Only pathways that were statistically significant at each applicable stage were retained.

To obtain a single precision-weighted causal estimate per miRNA, the standardized coefficients from Stage 1 and Stage 2 were combined using inverse-variance weighted (IVW) meta-analysis, with each stage weighted by the inverse of its squared standard error. Because lavaan does not directly report the standard error of the standardized coefficient, it was recovered as |β/z|, where z is the Wald statistic (the unstandardized estimate divided by its standard error), which is invariant to standardization. Prior to pooling, Stage 2 coefficients were sign-adjusted where canonical miRNA suppression produced opposing signs between stages, ensuring biological consistency of the pooled estimate. Relative to the side-by-side integration, this IVW pooling increases statistical power, reduces the combined standard error, and recovers pathways that act predominantly through the mediated (Stage 2) route. Therefore, the validated miRNA pathways were defined by the IVW meta-analysis.

### Causal inference using DoWhy

To validate and complement the SEM analysis, causal inference was performed using the Microsoft DoWhy library (v0.11; https://github.com/microsoft/dowhy). The underlying causal DAG was specified as shown in Fig. 1B, with age acting as a confounder simultaneously affecting miRNA expression, target gene expression, and AD status. The analysis proceeded in two steps: First, the causal effect of age on AD status (age→AD) was estimated using backdoor linear regression. The average treatment effect (ATE) was defined as the contrast between the 75th and 25th age percentiles (interquartile range, IQR contrast). Robustness was assessed using two refutation tests: a placebo treatment refuter (100 permutation simulations) and a data subset refuter (80% random subsets, 100 simulations). Second, the age→AD estimate was integrated with the SEM-derived standardized path coefficients from Stage 1 (miRNA → AD) and Stage 2 (target gene → AD) via IVW meta-analysis [37], Weights were defined as the inverse squared standard error of each stage estimate. This integration provided a precision-weighted causal summary across analytical layers, using age as an independently validated causal benchmark against which miRNA and target gene effect sizes were compared. Stage 3 (TF downstream target → AD) was not incorporated because it did not identify any additional miRNA pathways beyond Stages 1–2.

To evaluate whether TFs contributed additional causal signals in the exploration phase, a sensitivity analysis was performed. In this analysis, each miRNA’s Stage 2 latent factor was reconstructed from its combined target set (non-TF target genes together with its target TFs) rather than from target genes alone. If the combined target set did not support a latent factor, the most significant individual target was retained (observed-variable model). The age→AD estimate was integrated with this combined Stage 2 using the same IVW meta-analysis. In the validation phase, due to its smaller sample size, the integration used Stage 1 (miRNA → AD) and Stage 2 (target gene → AD) without TF incorporation or parceling. In this phase, the age→AD causal effect was estimated separately in each validation dataset and then pooled across cohorts by inverse-variance weighted meta-analysis (with between-cohort heterogeneity assessed via the I² statistic), yielding a single cross-cohort age benchmark. This cross-cohort pooling step was not required in the exploration phase, which was based on a single cohort. Only pathways demonstrating statistically significant effects across applicable stages were reported as validated causal pathways.

### Machine learning models for AD prediction

Based on causal analysis, several groups of miRNA candidates were chosen and compared for their predictive accuracy with several ML/DL models. TreeBagger, the random forest, SVM, and NN were adopted and compared. For details, refer to previously published articles [38, 39]. A clinical prediction model for AD was constructed through accuracy assessment, clinical effect assessment, and risk prediction, as shown in Fig. 1C. To further enhance performance, we applied a deep learning model, the Multi-Layer Perceptron (MLP) [40, 41], a fully connected feed-forward neural network capable of capturing non-linear feature interactions, offering a fundamentally different learning paradigm from the ensemble methods previously tested.

### Functional analysis

To characterize significant miRNA pathways from LSD analysis, several bioinformatics analyses were applied to determine the associations between those miRNA pathways and the clinical characteristics of AD samples from the GEO database, as shown in Fig. 1C. Gene Ontology (GO), Kyoto Encyclopedia of Genes and Genomes (KEGG) pathway, g:Profiler, Reactome (REAC), and WikiPathways (WP) analyses were performed on the significant pathway related genes between AD and controls.

### Data visualization and AI usage

Most plots were generated using R packages, while some were created manually or using the bioinformatics.com.cn online platform. The assembly and editing of the figures were performed using Inkscape (v.1.4.2), GIMP (v.3.2.4), and PowerPoint.

Claude Code (Anthropic) assisted in implementing the R and Python code required for portions of the data analysis and visualization. All AI-assisted code was reviewed, tested, and verified by the author against expected results. The methodological framework, including the design of the LSD pipeline (LASSO, SEM, DoWhy), method selection, parameter choices, model specifications, and interpretation of all results, were determined by the author. Additionally, editGPT (https://editgpt.app) was used to assist with language editing of the manuscript for grammar and clarity while preserving the author’s original writing style. The author takes full responsibility for the integrity and accuracy of all analyses and content.

### Statistics

All statistical analyses were conducted in R (v4.2.2) and Python (v3.9). For differential expressions of miRNAs across AD, MCI, and NC groups, the Jonckheere-Terpstra (JT) trend test was applied, and both p and q values were calculated for each miRNA. For differential gene expressions, Student’s t-test was used with p values (with two-tails) calculated for each gene. Odds ratios (OR) for miRNAs and TFs were estimated by univariate logistic regression, with AD status (AD = 1, NC/ND = 0) as the binary outcome, yielding ORs with 95% confidence intervals. For SEM analysis, the WLSMV and MLR estimators were applied in Stage 1 and Stage 2 and 3, respectively, and model fit was evaluated using three indices: CFI, RMSEA, and SRMR. For causal inference, the DoWhy library (Python) was used to estimate and validate the causal effects of miRNAs on AD status, using the backdoor criterion for identification and linear regression as the estimation method.

## Results

### Data Composition and Filtering of Target miRNAs and Genes

This study utilized the GSE120584 [21] and GSE33000 datasets in the discovery stage. GSE120584 contains miRNA profile data from blood samples of Alzheimer’s disease (AD) patients, mild cognitive impairment (MCI) patients, and normal controls (NC). GSE33000 [22] comprises mRNA profile data from brain tissue samples of AD and non-dementia (ND) controls. For validation, blood miRNA datasets of GSE215789 [23] and GSE46579 [24], and brain tissue mRNA datasets of GSE36980 [25] and GSE122063 [26]. were included in the stage analysis (details see Methods section).

Jonckheere-Terpstra (JT) trend analysis [42] of the miRNA characteristics among the AD, MCI, and NC groups in the discovery stage revealed 720 significant miRNAs (q<0.05), with 622 showing an increasing trend and 98 a decreasing trend from total 2562 miRNAs, indicating a more than six-fold greater increase than decrease. The statistical analysis is present in Table S1 of the Supplementary Materials. For mRNA GSE33000 dataset from brain tissue samples, Student’s t-test analysis of the mRNA profile between AD patients and ND controls revealed 27,022 significantly expressed mRNAs (p<0.05), as shown in Table S2, with 17,725 gene symbols and 9,297 accession numbers; those 17,725 genes were used for the following studies.

TargetScan (TS) [43] and miRDB were used for selecting miRNA candidates and their targets. The miRNA target gene candidates were assessed using multiple conservation scores, such as the context++ score of TS. By overlapping miRNAs with a context++ score of ≤ -0.50 in TS and a score of ≥ 80 in miRDB, we filtered 48 miRNA candidates, as shown in Fig. 2A. When using different context++ score thresholds of ≤ -0.25, -0.40, -0.45, and -0.50, the filtered miRNA candidates showed 51, 50, 48, and 48 with only one or two differences. Therefore, the context++ score of ≤ -0.50 from TS was selected for filtering miRNAs and their target genes.

**Figure 2.**
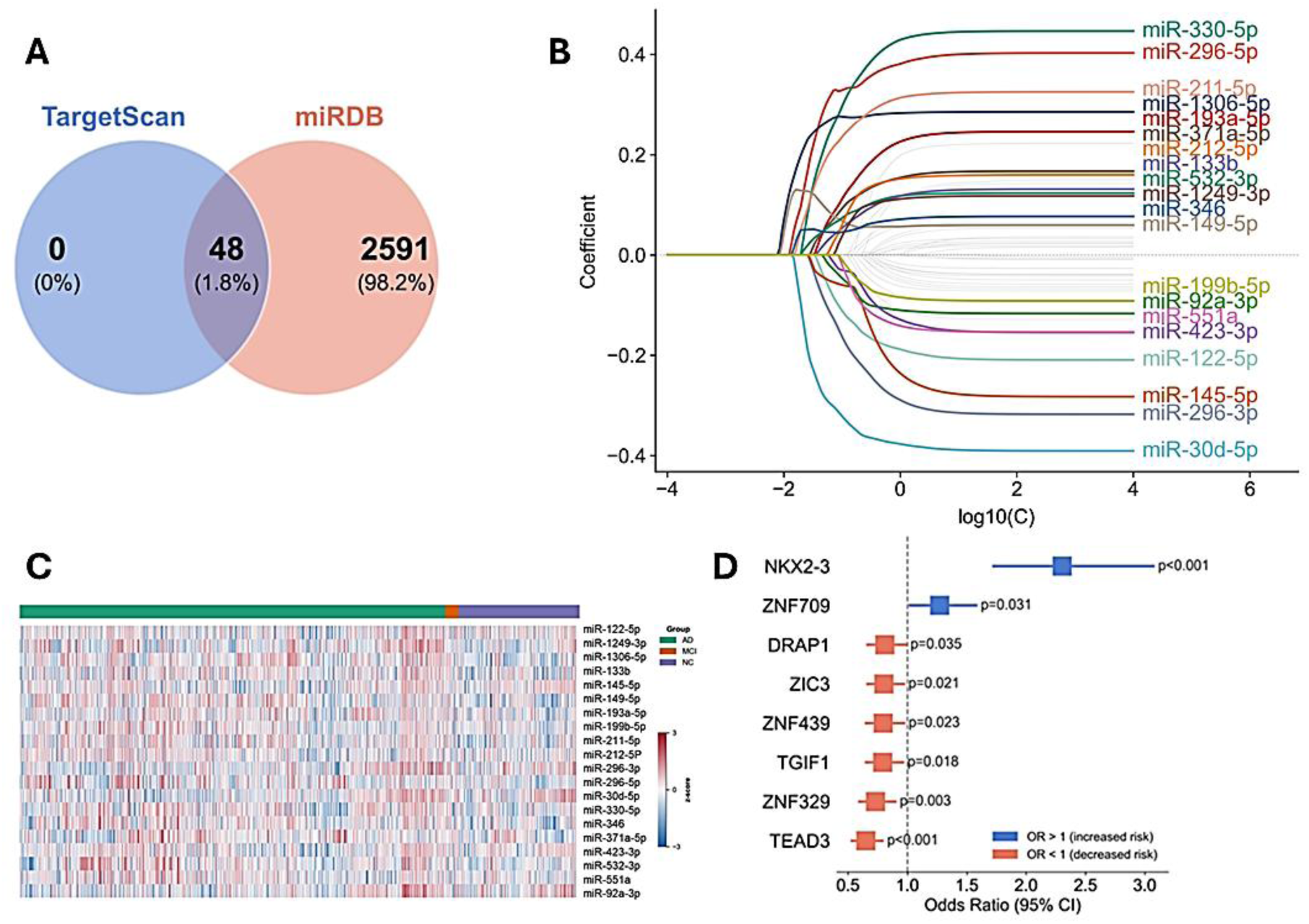
Pre-screening of miRNAs and Targeted TFs. A) 48 miRNAs were pre-filtered using the TargetScan and miRDB databases. B) LASSO logistic regression coefficient path for the 20 selected miRNAs based on the GSE120584 dataset. C) Expression heatmap of the 20 miRNAs selected by LASSO in GSE120584. D) Nine TFs were filtered and significantly associated with AD and their corresponding odds ratios based on the GSE33000 dataset. Red squares indicate an odds ratio < 1, suggesting a protective factor, while blue squares indicate an odds ratio > 1, indicating a risk factor.

### Pre-screening of miRNA Biomarker Candidates Based on LASSO Logistic Regression

To reduce the miRNA set to candidates for causal modeling, we applied LASSO logistic regression [38, 44]. We identified 20 significant variables (Fig. 2B) among 48 candidates, all of which were part of the pool of JT significant miRNAs (q<0.05). Univariate logistic regression (ULR) for each of the 20 miRNAs showed that 14 miRNAs increased the odds of AD, while 6 miRNAs decreased the odds, as illustrated in Fig. S1A. The expression levels of these miRNAs are presented in the heatmap (Fig. 2C).

Following the LASSO analysis, we filtered the target genes of the 20 miRNAs using the TS method and mapped them to the significant mRNAs. With a context++ score of ≤ -0.50, we identified 206 unique target genes that passed the significant filter (p<0.05) from the GSE33000 t-test analysis comparing AD and ND controls. Additionally, we established the relationships between the 20 miRNAs and their target genes, showing that some miRNAs regulate multiple targets (Table S3). Among the filtered target genes, we identified 192 non-transcription factors (non-TFs) and 14 transcription factors (TFs). To examine the co-expression relationships among the 14 TFs, the Pearson correlation matrix among the 14 TFs is shown in Fig. S1B. Several TF pairs showed statistically significant correlations (p < 0.05), suggesting coordinated regulatory activity within this network. For each TF, ULR results showed that nine of the 14 TFs were significantly associated with AD (p < 0.05), as shown in Fig. 2D. NKX2-3 showed the strongest positive association (OR = 2.30, p < 0.05); while TEAD3 showed the strongest inverse association (OR = 0.65, p < 0.05), suggesting its protectively regulatory role in AD pathology. Notably, the ZNF family members exhibited multiple significant inter-correlations, consistent with their shared structural class, e.g., ZNF439 and ZNF329 showed the strongest protective correlation (p < 0.05), while ZNF709 showed the positive correlation.

### Potential Causal Effects of AD by SEM Analysis

To investigate the causal relationships between serum miRNAs and AD, we applied two-stage or three-stage structural equation modeling (SEM) based on the 20 LASSO-selected miRNAs and their filtered target genes in the brain, sequentially linking serum miRNA levels → brain target gene expression → AD status. Model adequacy was confirmed by three complementary fit indices: comparative fit index (CFI), root mean square error of approximation (RMSEA), and standardized root mean square residual (SRMR), all of which indicated acceptable fit (Table S4).

Stage 1 SEM revealed that eight of the 20 miRNAs significantly predicted AD status (Table 1, Fig. 3A). In Stage 2, we examined whether all 20 miRNAs influence AD by modulating target gene expression in the brain (Fig. 3B). Table 1 shows six significant miRNA-related pathways before parceling, two of which do not overlap with those from Stage 1. After parceling, we identified ten significant miRNA pathways, including the initial six in Stage 2. Notably, five miRNAs did not converge in this stage. While Stage 1 β values [19] were predominantly positive, indicating that higher serum miRNA levels are associated with increased AD risk, the Stage 2 β values for the corresponding target genes were predominantly negative (Fig. 3B). This suggests that these miRNAs suppress their target gene expression. Since many of these targets are functionally protective, their reduced expression contributes to AD progression, consistent with the observed effects in Stage 1. In Stage 3, one TF target gene shows significantly positive effects on AD progression, while four TFs exhibit significantly protective effects, as shown in Fig. 3C. However, the interpretation is complex, as TFs can act as either transcriptional activators or repressors. Consequently, a single miRNA may upregulate some target genes while downregulating others, leading to mixed effects on AD progression. The three-stage SEM results are summarized in Table 1.

**Figure 3.**
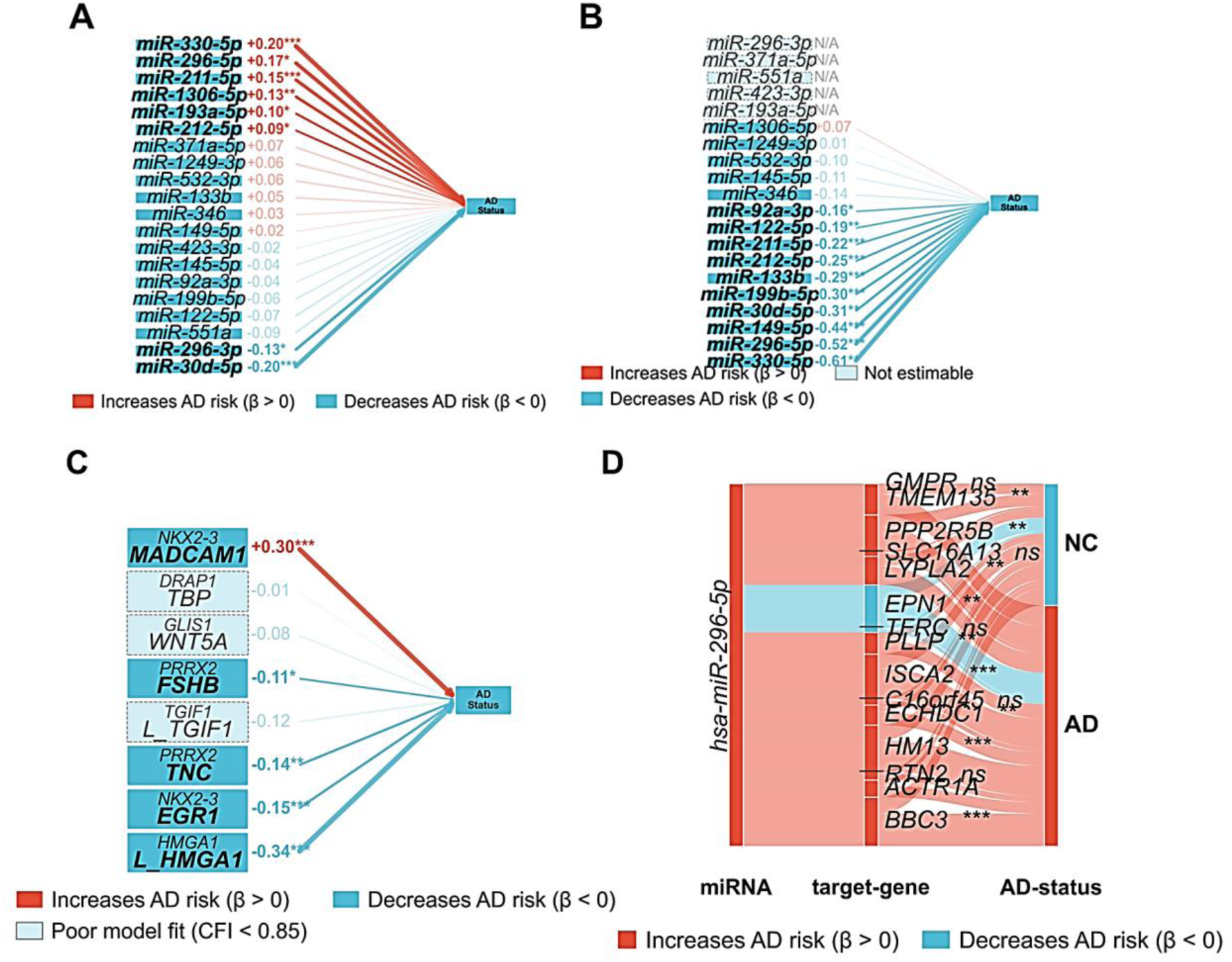
Three-Stage SEM Analysis and AD Causal Pathway Diagrams for miRNAs and Their Targets. A) Stage 1 of SEM analysis for 20 LASSO miRNAs to AD progression. B) Stage 2 of SEM analysis for all non-TF target genes of the 20 miRNAs to AD progression, including parceling analysis. C) Stage 3 of SEM analysis for all TF and their downstream target genes to AD progression, including the parcel process. D) Causal pathways of miR-296-5p for AD. Small black bars in the middle panel separate the parceling groups of target genes. In these plots, red lines indicate β > 0, signifying an increased AD risk, while blue lines indicate β < 0, signifying a decreased AD risk. *p< 0.05, **p <0.01, and ***p<0.001.

**Table 1.** miRNA–AD Path Significance across Three-stage SEM and DoWhy Analysis.

| miRNA | Stage 1 |  | Stage2 |  | Parceling |  | DoWhy |
| --- | --- | --- | --- | --- | --- | --- | --- |
|  | p-value | sig | p-value | sig | p-value | sig | sig |
| <u>miR-330-5p</u> | 0.000068 | *** | 1.29E-05 | (***) | 0.005746 | ** | *** |
| <u>miR-30d-5p</u> | 0.000191 | *** | 0.548964 |  | 0.007765 | ** | *** b |
| <u>miR-211-5p</u> | 0.000602 | *** | 3.10E-08 | *** |  | ** | *** |
| <u>miR-1306-5p</u> | 0.004739 | ** | 0.383047 |  |  |  | ** a |
| <u>miR-296-3p</u> | 0.015131 | * |  |  |  |  | * a |
| <u>miR-296-5p</u> | 0.031128 | * | 0.000504 | *** | 8.84E-07 | *** | *** b |
| <u>miR-193a-5p</u> | 0.031356 | * |  |  |  |  | * a b |
| <u>miR-212-5p</u> | 0.049039 | * | 0.010474 | * | 4.10E-07 | *** | *** |
| <u>miR-551a</u> | 0.120013 |  |  | ### |  |  | ### |
| <u>miR-122-5p</u> | 0.131431 |  | 0.153475 |  | 0.002758 | ** | ** |
| <u>miR-371a-5p</u> | 0.132281 |  |  |  |  |  |  |
| <u>miR-532-3p</u> | 0.247272 |  | 0.206905 |  |  |  |  |
| <u>miR-133b</u> | 0.268239 |  | 1.05E-09 | *** | 2.14E-10 | *** | *** |
| <u>miR-199b-5p</u> | 0.365678 |  | 9.41E-06 | *** | 6.46E-14 | *** | *** |
| <u>miR-1249-3p</u> | 0.367948 |  | 0.936362 |  |  |  |  |
| <u>miR-92a-3p</u> | 0.458645 |  | 0.608737 |  | 0.037399 | * | *** a b |
| <u>miR-346</u> | 0.507799 |  | 0.818544 |  |  |  |  |
| <u>miR-145-5p</u> | 0.516281 |  | 0.079876 |  |  |  | b |
| <u>miR-423-3p</u> | 0.743413 |  |  |  |  |  |  |
| <u>miR-149-5p</u> | 0.746191 |  | 0.161212 |  | 1.41E-05 | *** | * |

\*, \*\*, \*\*\*, p< 0.05, 0.01, or 0.001; <sup>a</sup> -items exist extra significant paths in DoWhy of exploration stage; <sup>b</sup> -items are significant miRNAs in validation stage; total 10 miRNAs exist in validation stage; ###, p<0.01 extra adding from TF stage2 and DoWhy analysis.

For individual miRNAs, the causal pathway of miR-296-5p is shown in Fig. 3D. For miR-296-5p, its 15 non-TF target genes underwent SEM analysis individually, after which they were divided into five parceling groups for further SEM analysis. After integrating, BBC3, HM13, and ISCA2 are the top three gene targets with highest loadings in the miR-296-5p-mediated pathway. Significant pathways are indicated with an asterisk (*), while increases or decreases in AD risk are represented by red or blue coloring, respectively. Among individual miRNAs, miR-133b and miR-199b-5p were not significant in Stage 1 (Table 1) but exhibited strong effects on AD in both Stage 2 and parceling analyses, indicating that their influence operates primarily at the gene-network level but is hidden at miRNA regulatory levels. These hidden miRNAs may not be detected by current research methods. In contrast, miR-296-3p, miR-193a-5p, and miR-1306-5p were predominantly involved in AD regulation only during Stage 1, not through their target genes, and may also be difficult to study currently.

Fourteen TF target genes of the 20 miRNAs were analyzed in stages 2 and 3 of SEM. In stage 2, the significant TFs included NKX2-3 and TGIF1 (miR-92a-3p), ZNF329/ZNF439/ZNF709 (miR-199b-5p), TEAD3 (miR-296-5p), and DRAP1 (miR-211-5p), while HMGA1 (miR-296-5p), PRRX2 (miR-212-5p), ZNF514 (miR-532-3p), and GLIS1 (miR-145-5p) were not significant. Three other TFs failed to converge. In stage 3, six TFs with eight downstream targets were evaluated (Fig. 3C). The results differed from those of stage 2. HMGA1 (β=-0.340, p<0.0001), previously non-significant in stage 2, became significant through its 55-gene downstream latent network after parceling. Similarly, PRRX2, non-significant in stage 2, exhibited significant downstream effects via TNC (β=-0.138, p<0.01) and FSHB (β=-0.108, p<0.05). NKX2-3 demonstrated further downstream effects via EGR1 (β=-0.146, p<0.001) and MADCAM1 (β=+0.305, p<0.001). Consequently, HMGA1 (High Mobility Group AT-Hook 1), PRRX2 (Paired Related Homeobox 2), and NKX2-3 (NK2 Homeobox 3) were identified as significant causal TFs of AD after SEM analysis. In contrast, TGIF1 was significant in stage 2, but it lost significance in stage 3, irrespective of whether its 10-gene downstream network was modeled as a single latent factor or through parceling. Additionally, DRAP1 and GLIS1 also lost significance in stage 3. Importantly, similar to the miRNAs, we revealed the "hidden network" effects of TFs that current research methods may not detect.

### Functional Analysis in Exploring Phase

To characterize the biological functions of SEM-identified miRNA target gene networks, functional analyses were conducted for the filtered target genes of four miRNA gene sets with the strongest multi-level SEM evidence: miR-199b-5p (24 non-TF targets), miR-133b (43 non-TF targets), miR-296-5p (15 non-TF targets), and miR-211-5p (10 non-TF targets). Additionally, the transcription factor HMGA1, which is one of the target genes of miR-296-5p, has 55 target sites identified in the 3-stage model and was also included.

The chord diagram (Fig. 4A) illustrates the AD-related pathways shared among the target genes of these four miRNAs. Several pathways converge on tau pathology through distinct mechanisms. For example, miR-296-5p→PPP2R5B→PP2A (GO:1904528) encodes the catalytic subunit of PP2A, the primary serine/threonine phosphatase responsible for tau dephosphorylation, while miR-133b→PP2CA/CB→PP2A (GO:1904528) further diversifies PP2A-tau interactions. Additionally, miR-199b-5p→MKMAP3K11→JNK signaling cascade (GO:0007254) promotes pathological tau hyperphosphorylation in AD.

**Figure 4.**
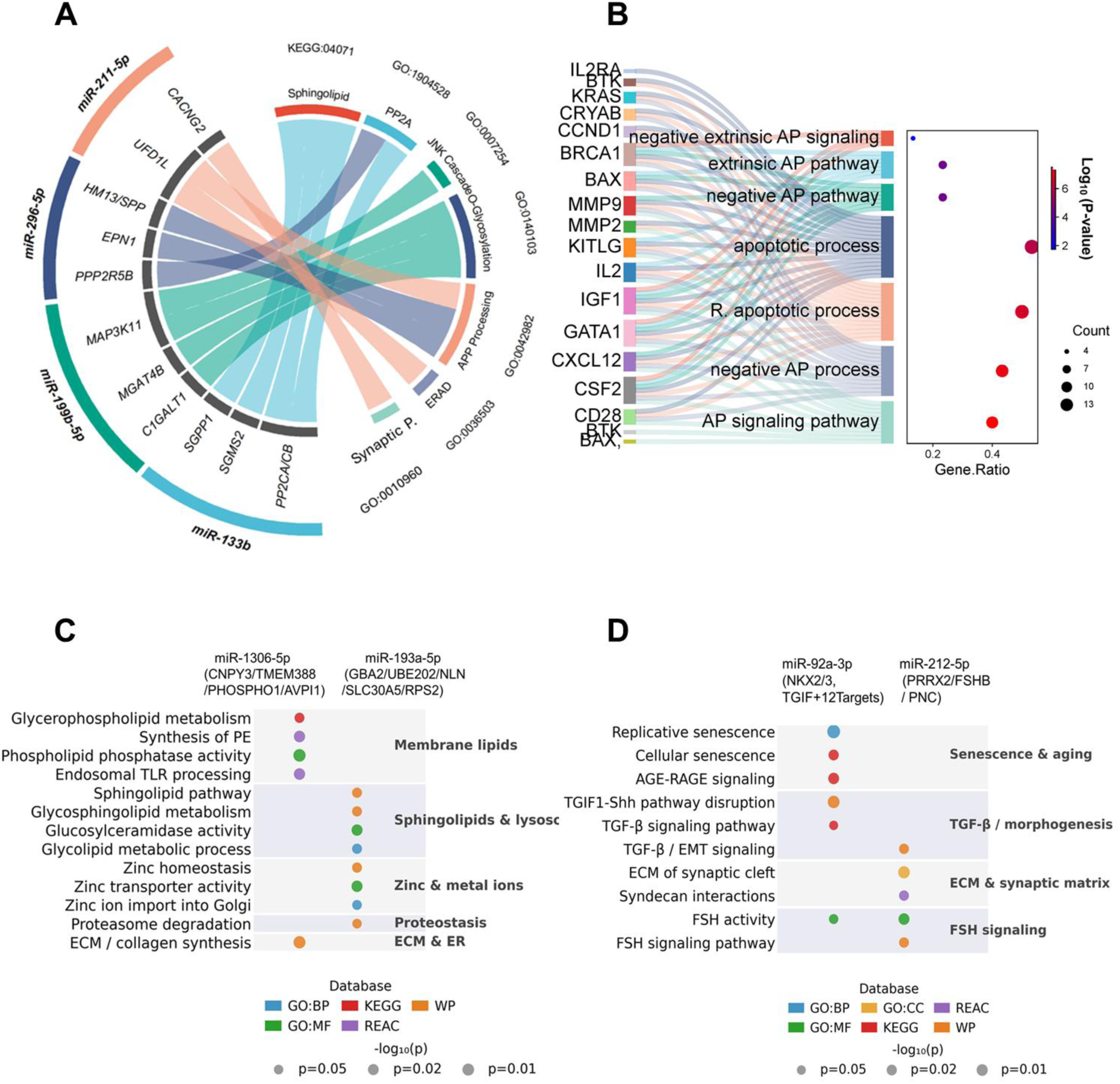
Function Analyses in the Discovery Phase. A) AD-related pathological pathways shared across miR-133b, miR-199b-5p, miR-296-5p, and miR-211-5p. From left to right, same color chords link a miRNA to its targets and GO terms in the chord diagram. B) Apoptotic enrichment pathways through TMGA1 downstream significantly expressed target genes. In the plot, R. and AP represent regulation and apoptotic, respectively. C) DoWhy analysis identified significant additional miRNA–non-TF pathways of AD and their functional analyses, including miR-1306-5p and miR-193a-5p. D) Functional analyses for miR-92a-3p and miR-212-5p related to TF pathways of AD.

HMGA1 is a TF target gene of miR-296-5p, which is used in the three-stage causal pathway analysis (miR-296-5p→HMGA1→its target genes→AD). Although it was not significant in stage 2 of the SEM analysis, it gained significance with its downstream targets in stage 3 with parceling. HMGA1 has 55 downstream target genes, 30 of which were differentially expressed genes (DEGs). Running g:Profiler with these DEGs reveals apoptotic enrichment pathways in Fig. 4B, from a total of 208 significant enrichment pathways. These apoptotic pathways encompass both the regulation of apoptosis and the negative regulation of apoptosis, which are directly related to AD. Additionally, three ZNF family members, ZNF329, ZNF439, and ZNF709, were identified as the target genes of miR-199b-5p. Importantly, none of these ZNF TFs currently have annotated downstream targets in curated databases, leaving their downstream regulatory programs unresolved and designating them as high-priority candidates for future experimental characterization.

To confirm the causal effects of miRNAs and their target genes through SEM analysis, DoWhy was performed, including age as a confounder, following the directed acyclic graph (DAG) structure outlined in Fig. 1B. In all DoWhy runs, the effect observed in the data subset was very close to the main average treatment effect (ATE) (Fig. S5). Additionally, placebo permutation tests yielded near-zero effects, confirming that age is a genuine and robust confounder rather than a spurious association. After the cross-stages in the DoWhy analysis for non-TFs as targets, three additional miRNAs reached significance: miR-1306-5p (**p < 0.01), miR-296-3p (*p < 0.05), and miR-193a-5p (*p < 0.05) (Table 1). These three pathways were not observed in the no-age SEM analysis in stage 2 for AD, where hidden network miRNAs were also identified. For the DoWhy analysis of miR-92a-3p, the results did change it was not significant in non-TFs analysis but gained significance in the TFs analysis. The combined gene+TF model was concordant with the gene-only analysis, preserving all primary miRNA pathways, and additionally recovered one miRNA: miR-551a reached significance in the combined Stage 2 and IVW meta-analysis through its target gene LPHN1 rather than a transcription factor. This underscores the value of the two-stage multi-omics design and cross-cohort integration, which are necessary for DoWhy confirmation and highlight the complexity of miRNA regulations.

Functional analysis was conducted for the three additional age-revealed miRNAs. For non-TF target genes (Fig. 4C), the targets of miR-1306-5p were enriched in phosphocholine phosphatase activity (GO:0052731), phosphoethanolamine phosphatase activity (GO:0052732), extracellular matrix (ECM) constituent secretion (GO:0070278), and synthesis of phosphatidylethanolamine (REAC:R-HSA-1483213). These findings indicate roles in membrane lipid homeostasis and innate immune signaling. The targets of miR-193a-5p were enriched in the sphingolipid pathway (WP:WP1422) and glycolipid metabolic processes (GO:0006664), implicating lysosomal lipid catabolism and proteostasis. For the related TF and TF target gene pathways of these three miRNAs (Fig. 4D), the results remain consistent regardless of whether age is considered a confounder. Functional enrichment analysis of miR-92a-3p targets (NKX2-3, TGIF1, and their downstream genes) revealed pathways associated with replicative senescence (GO:0090399) and cellular senescence (KEGG:04218), directly relevant to neurodegeneration in AD. Additionally, TGF-β signaling (KEGG:04350) and TGIF1-mediated Shh pathway disruption (WP:WP3674) illustrate the dual roles of NKX2-3 and TGIF1 in senescence and developmental signaling.

For the related TF and TF target gene pathways of these three miRNAs (Fig. 4D), the results remain consistent regardless of age as a confounder. Functional enrichment analysis of miR-92a-3p targets (NKX2-3, TGIF1, and their downstream genes) revealed pathways linked to replicative senescence (GO:0090399), cellular senescence (KEGG:04218), and AGE-RAGE signaling in diabetic complications (KEGG:04933), all relevant to neurodegeneration in AD.

For miR-212-5p, although not included in the DoWhy analysis, its TF target PRRX2 has two downstream targets: TNC and FSHB, which became significant at stage 3 of SEM, similar to HMGA1. These TFs function as ’hidden network’ TFs, analogous to ’hidden network’ miRNAs. The pathways enriched for the PRRX2 target and its downstream targets FSHB and TNC include the extracellular matrix of the synaptic cleft (GO:0098965), syndecan interactions (REAC:R-HSA-3000170), and TGF-β signaling in epithelial–mesenchymal transition (WP:WP3859) in Fig. 4D. The identification of hidden network TFs across both parceled (HMGA1 with 55 targets) and un-parceled (PRRX2 with 2 targets) stage 3 models of SEM suggest that target-aware causal modeling, independent of the indicator consolidation strategy, is the key analytical feature enabling their detection.

### Validation Results and Functional Analyses

For validation, two miRNA datasets from serum were utilized. GSE215789 is a sequencing dataset that detected 1,176 miRNAs across 112 samples, and GSE46579 is another sequencing dataset from serum, encompassing 503 miRNAs across 70 samples. After separately calculating z-scores for these two miRNA datasets, the combined dataset comprised 119 samples and included 10 miRNAs (underlined in Table 1) overlapping with the 20 LASSO-selected miRNAs, which were used for SEM and DoWhy analysis. Their expression heatmap was shown in Fig. S2. For mRNA validation, the GSE36980 dataset with 80 samples from frontal cortex (FC) tissue was combined with the GSE122063 dataset with 23 samples also from FC tissue, resulting in a total of 103 samples (45 AD and 58 controls). From the relationship data of the miRNAs and their target genes, 50 non-TF genes and two TF genes were identified in the combined dataset. The functional analysis of all 52 genes is presented in Fig. 5A. Significant GO terms included vesicle-related terms, cytosol, cytoplasm, and the endomembrane system, suggesting indirect regulation of AD progression.

**Figure 5.**
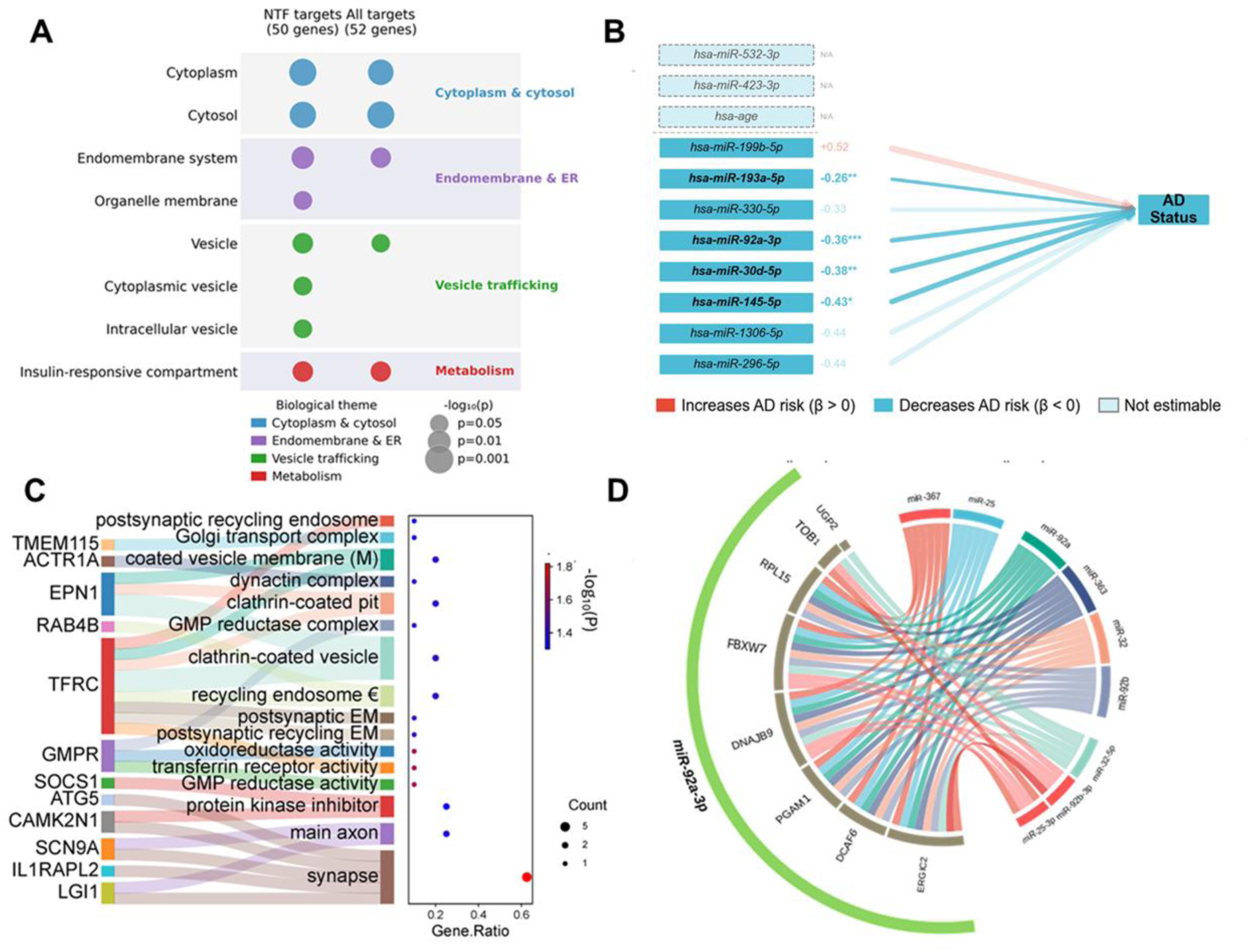
Validation Results and Their Functional Analyses. A) Functional analysis of all miRNAs related to 52 target genes. B) The pathway statuses of all miRNAs in Stage 2 of SEM analysis, highlighting four significant pathways. The blue color indicates a negative beta value. C) Significant AD-associated functional terms from miR-30d-5p and miR-296-5p. D) The Chord plot illustrates how the target genes of miR-92a-3p are linked to other miRNAs through functional terms.

Using the LSD package, SEM validation was performed with a two-stage causal model. A third stage for parceling was deemed unnecessary due to the limited number of targets, TF genes, and samples. In Stage 1, miR-30d-5p and miR-92a-3p were significantly associated with AD progression among the 10 miRNAs analyzed. In Stage 2, these two miRNAs remained significant, and miR-145-5p and miR-193a-5p also gained significance. Ultimately, four miRNA pathways (miR-145-5p, miR-193a-5p, miR-92a-3p, and miR-30d-5p) were significantly associated with AD progression, all exhibiting negative effects (β < 0), as shown in Fig. 5B, consistent with the exploratory stage. Neither TF gene (HMGA1 or TGIF1) demonstrated a significant effect, likely due to the small sample size (n = 23).

To assess the influence of age as a confounder, age-adjusted SEM was conducted using DoWhy causal inference within the LSD package. This analysis confirmed age as a significant causal factor for AD in both the miRNA (n = 119) and NTF gene cohorts (n = 103). After age adjustment, the Stage 1 results remained unchanged, with miR-30d-5p and miR-92a-3p being the only two significant miRNAs. However, in Stage 2, the set of four significant miRNA pathways remained: miR-145-5p, miR-193a-5p, miR-30d-5p, and miR-92a-3p persisted, and miR-296-5p gained significance. These changes enhance the consistency between Stage 1 and Stage 2, as both miR-30d-5p and miR-92a-3p are now significant in both stages. Additionally, after miR-145-5p was excluded due to its absence in the exploration phase, four key miRNAs (miR-30d-5p, miR-92a-3p, miR-193a-5p, and miR-296-5p) demonstrated consistency across the exploration and validation stages. Notably, these four key miRNAs are largely age-independent; miR-296-5p, for instance, was age-independent in the exploration phase, even though it gained significance after age was factored into the validation phase.

AD-associated GO terms for miR-30d-5p and miR-296-5p are presented in Fig. 5C. miR-30d-5p is associated with synapse (GO:0045202), main axon (GO:0044304), and protein kinase inhibitor activity (GO:0004860). The GO terms for miR-296-5p are primarily related to tau and amyloid-β pathologies of AD, as shown in the plot. GMP reductase activity (GO:0003920 and GO:1902560) is directly linked to AD, as overactivated kinases, including GSK-3β and MAPK, drive tau hyperphosphorylation and amyloid-β aggregation. Dysfunctions in endosomal recycling (GO:0098944 and GO:0055037) and membrane compartments (GO:0098895) lead to trafficking disruptions that contribute to hallmark AD pathologies, such as synaptic loss, and neuronal degradation. These functional terms are more specific than simply grouping every gene together (Fig. 5A), which is a standard process. For miR-92a-3p, no GO, KEGG, or Wiki pathway terms were identified due to the highly diverse functions of its target genes; however, several enriched miRNA terms were found, as shown in Fig. 5D. Since 2019, g:Profiler has included a miRNA category, and miR-92a-3p has been found to share regulatory networks with several other miRNAs, suggesting a broader upstream regulatory role of miRNAs in AD.

### Construction of ML Detection Model Based on miRNA Causal Effects on AD Analyses

To explore the potential clinical applications of miRNAs with causal linked to AD pathology, we combined the LASSO algorithm with the LSD causal analysis to identify candidate miRNA biomarkers and employed various ML/DL models for AD detection. In the exploration stage, we identified 20 important miRNAs through LASSO analysis from GSE120584, of which 10 overlapped with the validation datasets. Therefore, these 10 miRNAs were selected as a panel of biomarkers based on LASSO results. In the validation stage, four miRNAs (miR-30d-5p, miR-92a-3p, miR-296-5p, and miR-193a-5p) were identified as the key miRNAs from the LSD process, forming a second panel of biomarkers. Furthermore, due to its context-dependent significance, miR-193a-5p was excluded from the four-miRNA panel, resulting in a third panel comprising three miRNAs.

We initially utilized SVM, RF, and TreeBagger models [38, 39], with SVM achieving the highest accuracy among them in current studies. However, the AUCs of SVM model reached 0.714, 0.658, and 0.630 for the 10-miRNA, 4-miRNA, and 3-miRNA panels, respectively, using the GSE120584 dataset from the exploration phase (Fig 6A). For the combination dataset from the validation phase, the AUCs of SVM were 0.817, 0.810, and 0.758 for the same three panels, respectively (Fig 6B). To further enhance performance, we applied a deep learning model, the Multi-Layer Perceptron (MLP). With MLP, the AUCs for GSE120584 improved to 0.728, 0.670, and 0.629 for the same three panels, respectively (Fig 6C). For the combination dataset, the AUCs were 0.821, 0.861, and 0.762 for the same three panels (Fig 6D).The 4-miRNA panel yielded the highest AUC (above 86%) in the combination datasets of the MLP model analysis. These four miRNAs possess defined biological relevance and may hold potential for future clinical applications. All calculation details are shown in Table S5.

**Figure 6.**
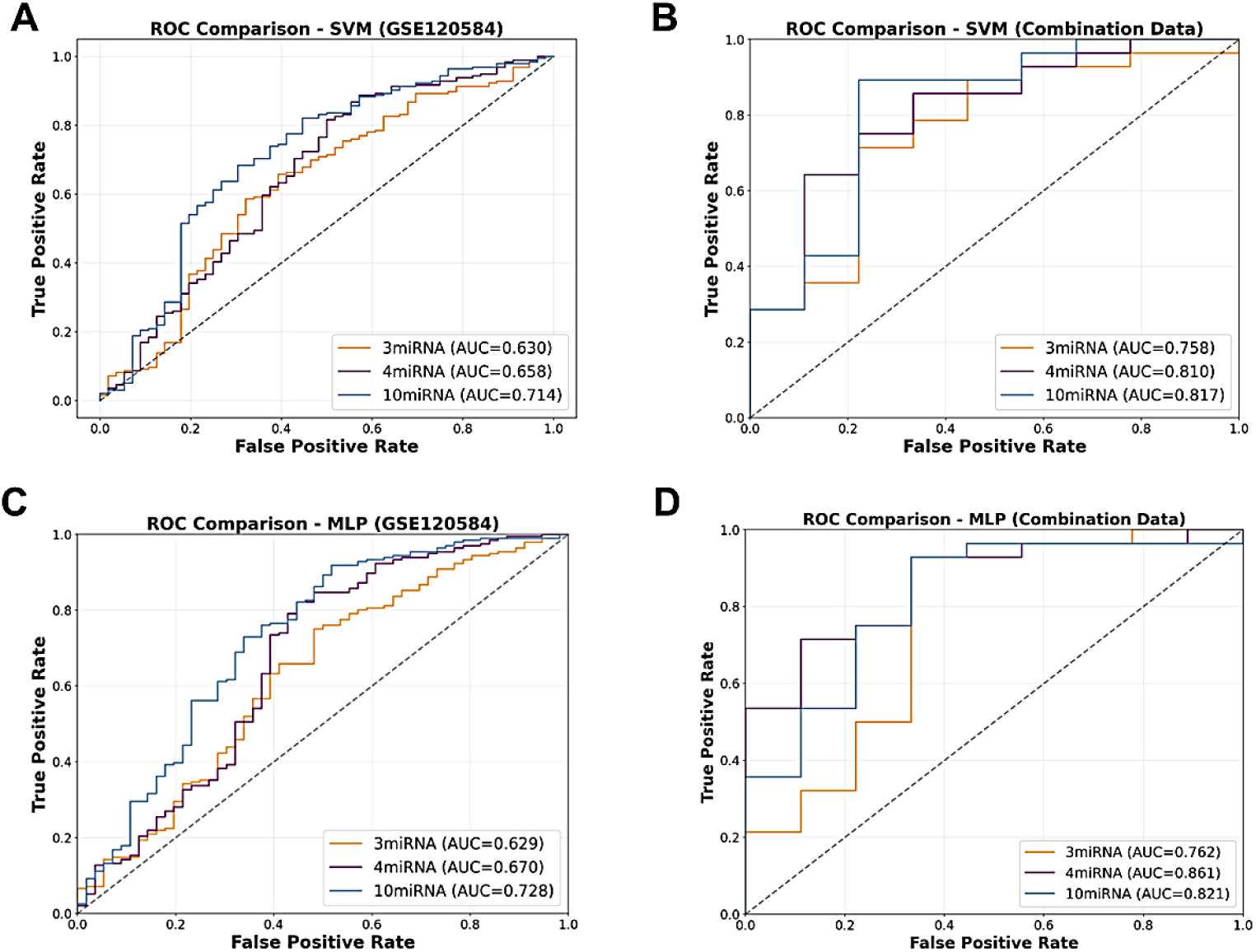
Construction of ML/DL Models Based on LASSO and LSD miRNAs for AD Diagnosis. A) ROC curves and AUCs of the SVM model detecting AD using three miRNA panels (3-miRNA, 4-miRNA, and 10-miRNA) in the exploration dataset (GSE120584). B) ROC curves and AUCs of the SVM model detecting AD using the three miRNA panels in the validation combination dataset. C) ROC curves and AUCs of the MLP model detecting AD using the three miRNA panels in the exploration dataset. D) ROC curves and AUCs of the MLP model detecting AD using the three miRNA panels in the validation combination dataset.

## Discussion

These results establish miRNAs as upstream causal regulators of AD pathogenesis, and they expose network-level mechanisms that conventional single-target analyses fail to capture. The LSD pipeline was developed precisely to address this gap.

### The LSD Pipeline

The LSD pipeline addresses two persistent challenges in miRNA research: the inability of conventional methods to model miRNA causal effects within regulatory networks and the reliance on statistical associations for biomarker selection without biological grounding. By integrating LASSO feature selection, multi-stage SEM, and DoWhy causal inference, LSD systematically elucidates miRNA causal pathways from blood-circulating miRNAs through brain mRNAs and TF cascades to AD outcomes. Unlike analyses limited to a single cohort or tissue type, our cross-cohort and cross-tissue design leverages the conserved binding specificity of miRNAs, enabling robust inference across diverse data sources.

A prior study in amyotrophic lateral sclerosis identified causal associations between miRNA-related genetic variants and disease using Mendelian randomization with miRNA-eQTL data [45], modeling miRNA expression as a mediator between genetic variants and disease. In contrast, LSD addresses a complementary question by modeling miRNA expression itself as causal exposure, capturing miRNA-driven mechanisms regardless of whether expression changes stem from genetic, environmental, or epigenetic sources. The pipeline confirmed classic AD pathological hallmarks, such as tau pathology and synaptic dysfunction, while also identifying novel TF candidates and an LSD miRNA panel with high predictive accuracy—biomarkers that possess explicit causal grounding rather than being selected purely on statistical grounds.

The consistent directional trends observed across all three LSD stages are predominantly upregulated miRNAs (β > 0), corresponding downregulated mRNA targets including protective factors (β < 0), and coherent TF cascade directions, which support LSD as a framework for exploring biologically meaningful regulatory architecture. Combined with the identification of age-independent and "hidden network" miRNAs and TFs across two independent cohorts, along with consistency with prior genetic and functional evidence, these findings suggest that LSD captures real network-level mechanisms rather than analytical artifacts.

### Age-Independence of miRNA Causal Effects

The causal effects of AD-associated miRNAs identified by the LSD pipeline were largely independent of age, while the downstream mRNA and TF cascades were highly sensitive to age. This pattern remained consistent across both exploratory and validation datasets. Given that age is a primary risk factor for AD, this finding is unexpected and has not been previously reported. However, it has been noted that grief-related stressful life events (SLEs) are associated with cerebrospinal fluid (CSF) markers of AD, including Aβ42/40 and p-tau181, and that these associations are independent of age [46]. While that study attributes the age independence of the biomarkers to SLEs as the driving factor, our findings suggest a different mechanism.

We propose that the observed age independence reflects our initial filtering step: by applying stringent TargetScan context++ thresholds (≤ −0.50) [29, 43], we focused our analysis on miRNA-target pairs with highly conserved binding sites across species. Evolutionarily conserved miRNAs tend to regulate fundamental biological processes rather than age-specific pathways, which likely explain their stable expression across age groups, despite significant differences between AD patients and controls. In contrast, less stringent databases retained dozens of candidate targets per miRNA (up to 70 per miRNA), overwhelming both LASSO feature selection and SEM parameter estimation in current datasets. TargetScan’s conservation-based filtering reduced target sets to manageable sizes, enabling stable LASSO convergence and adequately powered SEM analysis. The age independence of LSD miRNAs has important clinical implications: these miRNAs may serve as biomarkers for the early prediction and detection of AD and may be linked to initial mechanisms of disease pathogenesis preceding age-related decline.

### Network-Level miRNA Effects in AD

Our findings highlight that miRNAs affect AD primarily through complex regulatory networks rather than unique individual miRNA-target interactions, as most previous studies have assumed. The LSD analysis revealed at least three distinct patterns of network behavior. First, certain miRNAs (miR-296-3p and miR-1306-5p) exhibited significant direct effects on AD that attenuated once their downstream target genes were modeled; yet they remained significant in the DoWhy causal analysis. The partial attenuation indicates that the modeled targets account for much of their influence, while the residual effect most likely reflects further mediating targets excluded by our deliberately conservative selection, rather than genuine direct action on AD. The LSD pipeline further models age, the dominant driver of AD, as a confounder, so the surviving effects reflect disease-specific regulation independent of aging, which analyses that omit age cannot isolate.

Second, other miRNAs (miR-133b, miR-199b-5p, and miR-149-5p) became significant only after modeling their target networks, as shown in Table 1. These "hidden network" miRNAs do not appear to regulate AD directly; instead, they exert their influence entirely through downstream targets, an effect that single miRNA-axis studies would overlook. This pattern is consistent with the post-transcriptional function of miRNAs: a miRNA acts upstream of its functional effects, which are realized in the coordinated response of its target genes rather than in its own expression level. Its abundance is therefore an incomplete proxy for its functional activity, and target-aware modeling recovers this latent regulatory signal. Its persistence after causal adjustment (DoWhy) indicates a genuine target-mediated route to AD rather than a modeling artifact.

Finally, miR-92a-3p showed context-dependent significance: it was significant in the stage-2 parceling analysis of the exploration cohort and confirmed by DoWhy, and it reached significance in the stage-2 without parceling in the validation cohort. Its target genes span functionally diverse pathways, so pooled GO and KEGG enrichment cannot resolve a coherent functional theme. Rather than paradoxical, this supports our central argument: miRNA regulatory function is multimodal and intrinsically dependent on the network context in which it operates. For such miRNAs, pooled enrichment is the wrong lens; examining individual miRNA–target pairs, the traditional approach, better reveals the distinct target roles that pooling obscures. This complements rather than replaces the network-level causal modeling, which detected the effect that enrichment alone could not interpret.

The miRNA network is more complex than these three patterns suggest: many miRNA targets are themselves transcription factors, adding regulatory layers that amplify network-level impact. These TF cascades are discussed next.

### TF Cascades and Key Regulators

Building on our previous findings that transcription factors (TFs) are involved in miRNA regulatory pathways in glioblastoma [38], we emphasized TFs in this study. We separated TFs from non-TF genes, modeling TF-mediated cascades as Stage 3 of the SEM analysis (Fig. 1). Our results indicate that TFs interact with miRNA regulatory networks through at least two distinct mechanisms.

The first mechanism involves TFs as direct miRNA targets: miRNAs repress TF expression, and the affected TFs regulate their own downstream genes. Since TFs can act as activators or repressors, they may drive AD progression in either direction even when suppressed by miRNAs. This miRNA→TF→downstream gene→AD cascade was confirmed for HMGA1, NKX2-3, and PRRX2 in Stage 3 SEM. Although prior studies have examined miRNA→ mRNA→TF networks in dilated cardiomyopathy [47] and gastric cancer [48], these typically identified TFs among DEGs and prioritized candidates through topology-based metrics without investigating downstream causal effects.

HMGA1 emerged as a major contributor to tau pathology. As a target of miR-296-5p with 55 downstream sites in the three-stage SEM model, HMGA1 expression was significantly reduced in AD patients across both exploratory and validation datasets. Prior work has linked HMGA1 deficiency to tau accumulation and behavioral abnormalities in tau transgenic mice and tauopathy patients [49], with the rs146052672 variant reducing HMGA1 expression and associating with lower MMSE scores [50]. In our analysis, HMGA1 was not significant in SEM without parceling but became significant with parceling (β = −0.34, p < 0.01; Fig. 3C), supporting a protective role whose loss contributes to AD risk. Validation analysis was limited by insufficient HMGA1-expressing cases (n = 23).

This pattern revealed ’hidden network TFs’, regulators significance only when downstream targets are incorporated. HMGA1 (55 targets) and PRRX2 (two targets, FSHB and TNC, from miR-212-5p) became significant in Stage 3 but not Stage 2. Conversely, TGIF1, DRAP1, and GLIS1 were significant in Stage 2 but lost significance with downstream modeling. Both patterns demonstrate that target-aware modeling is essential for accurate TF assessment.

The second mechanism involves TFs regulating miRNA targets from upstream (miRNA → target ← TF), without being miRNA targets themselves. Several ZNF-family TFs (ZNF709, ZNF329, ZNF439) emerged as significant causal nodes in miR-199b-5p pathways (Fig. 2D, Fig. S1B) but could not be evaluated by SEM due to unannotated downstream targets. ZNF439 has recently been associated with AD cortical tissue by TWAS [51], consistent with our findings, though its functional role remains to be determined experimentally. Additionally, some TF downstream targets are themselves TFs, suggesting multi-level regulatory cascades in AD pathogenesis that the current 3-stage LSD pipeline partially captures but does not fully model. Future extensions of this pipeline to incorporate recursive miRNA-driven TF–TF regulatory interactions may reveal additional causal pathways in AD and other complex diseases.

### Limitations and Future Direction

Several limitations should be acknowledged. As a computational study based on observational multi-omics data, the causal effects identified by the LSD pipeline depend on assumptions intrinsic to SEM and DoWhy. Experimental validation through RT-qPCR [52] and functional assays will strengthen these findings, although network-level mechanisms (such as miRNAs or TFs that act entirely through downstream cascades or "hidden network" regulators lacking prior experimental support) may require validation strategies tailored to network-level causality. The cross-tissue design infers pathways from blood miRNAs to brain mRNAs across separate cohorts and relies on conserved miRNA binding specificity across tissues. While supported by external validation, future studies with matched blood and brain samples would provide more direct confirmation.

The discovery and validation phases utilized different platforms (microarray vs. RNA-seq), which vary in miRNA coverage (>2,000 vs. >500). Consequently, miR-133b was identified and confirmed during the discovery phase but was undetected in the validation dataset, despite its reported presence in AD serum and tissue samples and its implication in synaptic integrity [53–55]. Despite these differences, LSD produced consistent causal effects across both platforms. Finally, only age was modeled as a confounder; the APOE genotype was unavailable for the validation datasets, and sex was excluded because its effects are mediated through hormonal pathways not captured by the available data. Future analyses will address APOE status, sex with hormonal information, and other AD risk factors.

Validation will require larger and more diverse AD cohorts, along with wet lab confirmation by RT-qPCR on tissue and blood samples. Investigating network-level mechanisms, particularly hidden-network miRNAs and TF cascades will necessitate experimental designs focused on network-level rather than single-target causality. Finally, applying LSD to other neurodegenerative diseases and cancers will help assess its generalizability and may reveal shared regulatory principles across miRNA-driven disease mechanisms.

Our study provides evidence that causal effects, bioinformatics, and AI studies in life sciences can uncover novel factors and principles beyond current experimental research. This approach is not only necessary but also more important than previously realized. It creates tools for pre-screening wet experiments and enables in-depth exploration of multi-omics, multi-tissue, and network-level relationships and associations. In fact, we currently know very little about regulatory networks involving multiple levels and components. Therefore, we need to further develop and explore basic principles using and combining this approach with experiments in various ways.

Conclusion: We developed the LSD (LASSO–SEM–DoWhy) pipeline to identify causal miRNA pathways in AD. We validated a four-miRNA panel (miR-30d-5p, miR-92a-3p, miR-296-5p, and miR-193a-5p) that demonstrated over 86% detection accuracy in the combined validation cohort using an MLP model. Our analysis uncovered not only individual biomarkers but also network-level behaviors of miRNAs and TFs: including "hidden network" regulators that operate through downstream cascades. We confirmed causal chains converging on tau pathology and synaptic dysfunction. By tracing complete causal pathways from miRNA to mRNA to TF cascades in AD, we identified HMGA1, NKX2-3, and PRRX2 as key intermediaries. The LSD pipeline enhances our understanding of miRNA-driven mechanisms in AD and provides a transferable framework for network-level causal inference in other diseases.

## List of abbreviations

AD: Alzheimer’s disease
LASSO: least absolute shrinkage and selection operator
LSD: LASSO–SEM–DoWhy
miRNA: microRNA
SEM: structure equation model
AET: average effect treatment
TF: transcriptional factor
HMGA1: high mobility group AT-hook 1
NKX2-3: NK2 Homeobox 3
PRRX2: paired related homeobox 2
TGIF1: TGFB-induced factor homeobox 1
DRAP1: DR1-Associated Protein 1
DAG: directed acyclic graph
GLIS1: GLIS family zinc finger 1
EGR1: early growth response 1
PPP2R5B: regulatory subunit for the PP2A
CFI: comparative fit index
RMSEA: root mean square error of approximation
MR: Mendelian randomization
KEGG: Kyoto encyclopedia of genes and genomes
GO: gene ontology
TWAS: transcriptome-wide Association studies
FSHB: follicle-stimulating hormone β
TNC: tenascin C
SLEs: grief-related stressful life events
DEG: differential expressed gene
MLP: multi-layer perceptron
MMSE: mini-mental state examination
SVM: support vector machine
RF: random forest
AUC: area under curve
ROC: receiver-operating characteristic
BBC3: BCL2 Binding Component 3
HM13: histocompatibility minor 1
ISCA2: iron-sulfur cluster assembly 2
SRMR: standardized root mean square residual

## Availability of data and materials

All datasets analyzed in this study are publicly available from the Gene Expression Omnibus (GEO; https://www.ncbi.nlm.nih.gov/geo/). The exploration analysis used blood miRNA data from GSE120584 and brain mRNA data from GSE33000. For validation, blood miRNA data from GSE215789 and GSE46579 were combined, and brain mRNA data from GSE36980 and GSE122063 were combined. No new data was generated in this study.

We generated the lsd_pineline for this study. The code is maintained at the UCSF GitHub repository (https://git.ucsf.edu/jian-shi/lsd_pipeline). The LSD pipeline (LASSO–SEM–DoWhy) is publicly available on Zenodo (https://doi.org/10.5281/zenodo.20369677) under the MIT License. The deposit contains source code in R and Python, a demo dataset, installation instructions with dependency versions, and scripts to reproduce all main analyses.

## Competing interests

The authors declare that they have no competing interests.

## Funding

This research was conducted without external funding.

## AI-assisted technologies

Claude Code (Anthropic) assisted in implementing the R and Python code used for portions of data analysis and visualization. All AI-assisted code was reviewed, tested, and verified by the author against expected results. Additionally, editGPT (https://editgpt.app) assisted with language editing of the manuscript for grammar and clarity while preserving the author’s original writing style. The author takes full responsibility for the integrity and accuracy of all analyses and content.

## Notes

https://www.ncbi.nlm.nih.gov/geo/query/acc.cgi?acc=GSE120584

https://www.ncbi.nlm.nih.gov/geo/query/acc.cgi?acc=GSE33000

https://www.ncbi.nlm.nih.gov/geo/query/acc.cgi?acc=GSE215789

https://www.ncbi.nlm.nih.gov/geo/query/acc.cgi?acc=GSE46579

https://www.ncbi.nlm.nih.gov/geo/query/acc.cgi?acc=GSE36980

https://www.ncbi.nlm.nih.gov/geo/query/acc.cgi?acc=GSE122063

https://doi.org/10.5281/zenodo.20369677

### Competing Interest Statement

The authors have declared no competing interest.

