## Supplemental Figures FigS1 and FigS2 for "LSD-pipeline: Causal Inference of miRNA Network Effects in Alzheimer’s Disease"

**Supplemental Materials**

**Title: Network-Level Causal Effects of miRNAs in Alzheimer's Disease**

**Fig. S1**

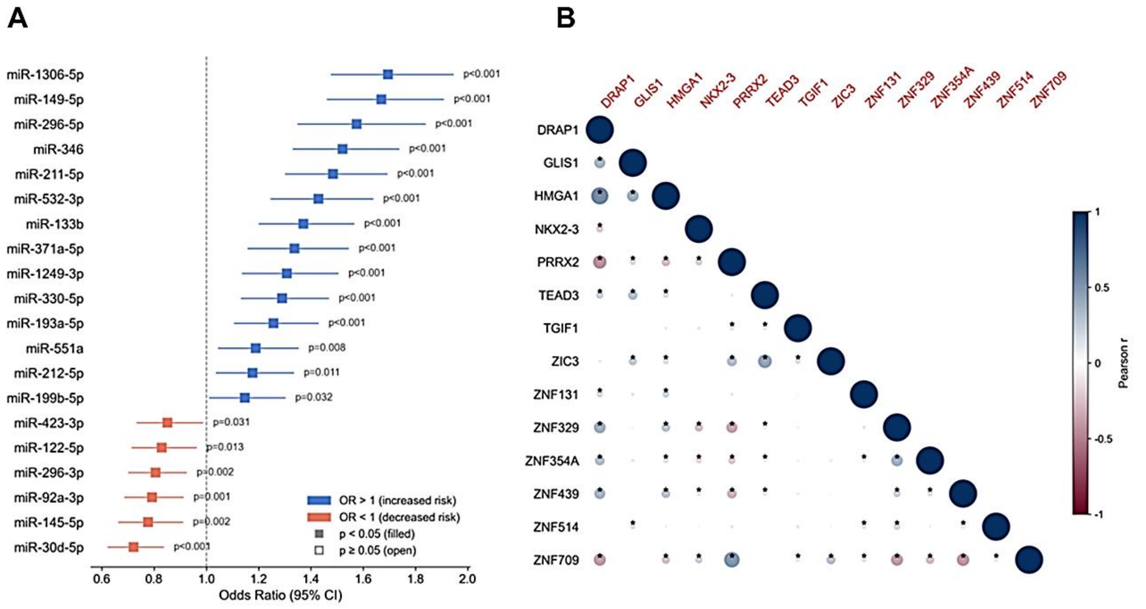


**Figure S1.** The AD odds ratios of miRNAs and the co-expression relationships of TFs. A) Among the 20 miRNAs selected by LASSO, a univariate logistic regression analysis revealed that 14 miRNAs increased the odds of AD ($p < 0.05$), while 6 miRNAs decreased them ($p<0.05$). B) The co-expression relationships among the 14 TFs were analyzed using Pearson correlation. Several TF pairs exhibited statistically significant correlations ($p < 0.05$), indicating coordinated regulatory activity within this network.

**Fig. S2**

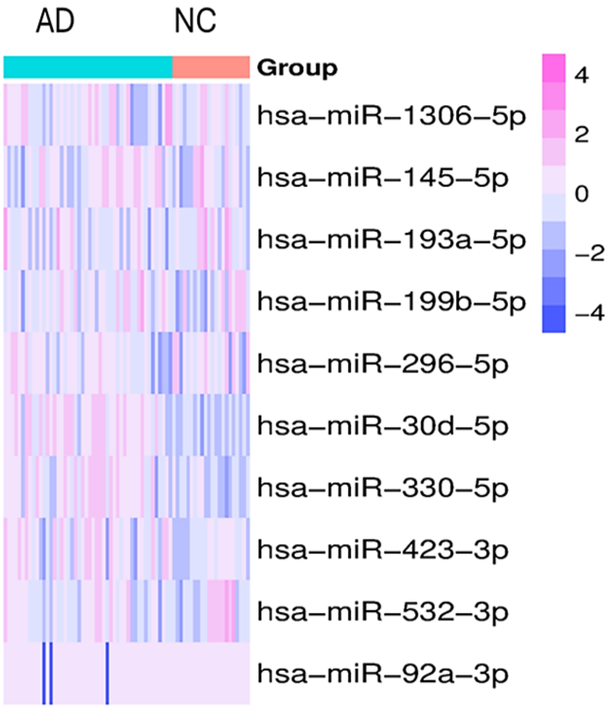


**Figure S2.** The heatmap for 10 miRNAs in validation phase. 10 miRNAs (underlined in Table 1) overlapped with the 20 LASSO-selected miRNAs, including miR-1306-5p, miR-145-5p, miR-193a-5p, miR-199b-5p, miR-296-5p, miR-30d-5p, miR-330-5p, miR-423-3p, miR-532-3p, and miR-92a-3p. which were used for SEM and DoWhy analysis.
